# A Generalization of the Ternary Binding Model to Membrane-Confined Systems with Finite Copy Number

**DOI:** 10.64898/2026.04.10.717668

**Authors:** Hamid Bellout, Angela Li, Konstantin Piatkov, Dean Bottino

## Abstract

Bispecific T cell engagers and related immunotherapies are dosed using equilibrium binding models derived for well-mixed solution, yet therapeutic activity occurs at nanoscale membrane synapses with finite receptor copy numbers. Here we show that membrane confinement introduces geometry-dependent corrections to the landmark Douglass [6] ternary binding model, shifting formation half-points (*TF*_50_) by 2–10-fold at clinically relevant antigen densities. We present two complementary formulations—effective concentration and surface density—that preserve the Douglass framework while explicitly accounting for synapse geometry, surface topology, and the *accessibility factor* (*η*) of surface receptors. We further derive stochastic descriptions of trimer formation via the chemical master equation, demonstrating the recovery of the classical Ternary Binding Model equilibrium in appropriate limits.

We illustrate the framework using the CD19-targeting BiTE blinatumomab as a case study. Accounting for microvillus-driven patchy close contact during immune surveillance yields a mechanistic explanation for why higher target antigen density can *increase* the dose required to achieve a fixed level of ternary formation: in the membrane-confined regime, excess target acts as a local *antigen sink* that sequesters drug and reduces the free fraction available for productive bridging. Rather than fitting to a single shift value, we emphasize the robust *scaling* and *regime structure* predicted by the theory (density-proportional behavior in the sink-dominated limit, and collapse toward affinity-limited behavior outside that limit). The generalized framework provides ready-to-use correction formulas and parameter-estimation guidance, establishing a rigorous foundation for antigen-density-aware dosing strategies in T cell engager pharmacology.

**Statement of Significance:** Bispecific T cell engagers are currently interpreted largely through bulk-solution binding models, but at nanometer-scale immune synapses those models can miss a distinct, decision-relevant regime. We generalize the standard ternary binding model to membrane-confined synapses and relate whole-cell receptor counts to effective concentrations in the contact zone. In a blinatumomab case study, the framework explains a paradoxical observation: increasing target density can shift half-maximal ternary-complex formation to higher doses because abundant antigen acts as a local sink that sequesters drug. These results show when bulk affinity alone is insufficient for potency interpretation and support antigen-density-aware dosing and experiments that distinguish geometric from chemical control.

## 1 Introduction

Many modern immunotherapies rely on a bridging ligand that simultaneously engages two distinct receptors, such as a tumor-associated antigen (TAA) on a cancer cell and the CD3 complex on a T cell. Bispecific T cell engagers (BiTE™), chimeric antigen receptor constructs, and related molecules all share this basic architecture. A central quantity of interest is the equilibrium or steady-state concentration of the productive trimeric complex formed between the two receptors and the bridging ligand, and how this concentration varies with ligand dose and receptor abundance.

Douglass and coauthors developed a comprehensive thermodynamic model for three-body binding equilibria in well-mixed, three-dimensional solution [6]. Their analysis revealed that, for a broad class of systems, the trimer concentration as a function of the total bridging ligand concentration is bell-shaped, with an initial formation limb, a maximal point, and an inhibitory limb at high ligand associated with sequestration into dimeric complexes. They introduced formation and inhibition half points, denoted *TF*_50_ and *TI*_50_, as useful reduced descriptors of this nonmonotonic response. We will refer to their model as the Ternary Binding Model (**TBM**).

In cell-based applications, however, the assumptions of the original model are violated in several important ways. First, the receptors are confined to opposed membranes and can only interact within a finite contact zone of area *A*_c_ and gap height *h*. Second, the relevant receptors are present in finite copy numbers per cell. Whole-cell counts are typically 10^4^–10^5^ molecules, of which a fraction *η* ∼ 10^−3^–10^−1^ is synapse-engaged, yielding 10^2^–10^4^ engaged molecules in the contact zone (Section 3.1). Third, bond formation and rupture are stochastic events, and the number of trimers present in a given synapse is an integer random variable, particularly when copy numbers are modest. Fourth, and critically, the actual reactive surface is determined not by the projected contact area but by the topology of membrane protrusions—microvilli—that create the initial contact points between cells.

These features motivate a refinement of ternary complex theory that respects membrane geometry, microvillus topology, finite copy number, and stochasticity while remaining anchored to the (**TBM**) equilibrium in appropriate limits.

In this work we construct such a refinement. We first recall the classical ternary equilibrium in bulk. We then present two membrane-confined generalizations. The first (Branch A) introduces effective three-dimensional concentrations for membrane-anchored receptors in the contact volume and directly substitutes these into the (**TBM**) algebra while tracking finite copy numbers explicitly. The second (Branch B) reformulates the problem entirely in terms of surface densities and two-dimensional dissociation constants. For both branches, we introduce a stochastic description of dimer and trimer counts and show how, in the long-time and large-copy limit, the deterministic (**TBM**) solution is recovered. Finally, we analyze the dependence of the formation half point *TF*_50_ on target antigen copy number and explain why it can shift to higher ligand doses as target density increases in sequestration-dominated regimes.

### Defining the Core Problem: From Encounter to Engagement

*To construct a truly predictive framework for therapeutic potency, we must focus on the gatekeeping phase of the cellular interaction, not only its endpoint. The formation of a mature synapse is the consummation of the process—effectively the stage where the game is over and close apposition makes adhesion strongly favorable. The determinant of potency is therefore whether the system can reach that point at all*.

*Our analysis isolates this critical, decisive phase: the transition from the loose scanning state to stable engagement. In this regime, cells are “close but not there yet,” separated by steric and integrin constraints (an intermembrane separation h on the order of* ∼ 50 *nm, rather than the* ∼ 15 *nm spacing of a mature synapse). While the Douglass framework is a chemical equilibrium description, we apply it here as a quasi-equilibrium nucleation criterion, computing the equilibrium abundance (and corresponding formation probability) of “seed” ternary complexes conditional on the scanning geometry. By quantifying this conditional bridging likelihood, we capture the will it or won’t it threshold that governs efficacy*.

*Modeling the mature synapse would describe the consequences of success; modeling the scanning regime predicts the likelihood of success*.

## 2 Bulk ternary equilibrium: summary of the Douglass model

We briefly summarize the key features of the ternary equilibrium model of Douglass [6]. Consider three molecular species A, B, and C in a well-mixed volume, with total concentrations [*A*]_*t*_, [*B*]_*t*_, and [*C*]_*t*_. Species B can bind A and C to form dimers AB and BC with dissociation constants *K*_*AB*_ and *K*_*BC*_, and the dimers can further associate to form a trimer ABC with dissociation constant *K*_*T*_ . The basic reactions are

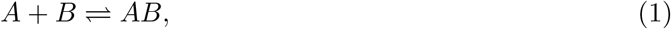

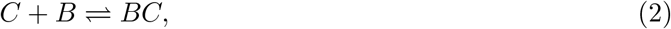

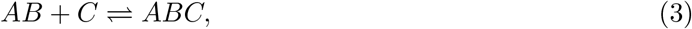

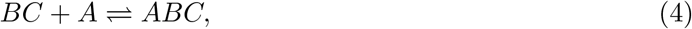

with associated equilibrium relations

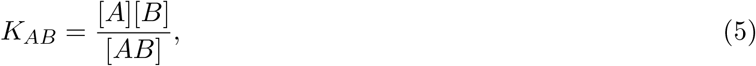

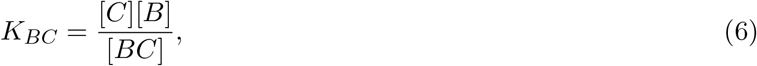

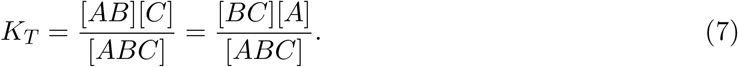

Mass balance gives

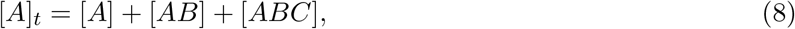

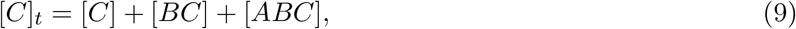

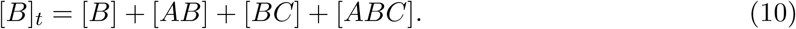

Douglass and coauthors expressed the dimer concentrations as quadratic functions of [*B*]_*t*_,

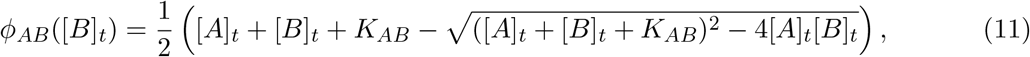

and analogously for *ϕ*_*BC*_([*B*]_*t*_) with [*C*]_*t*_ and *K*_*BC*_. The trimer concentration [*ABC*] can be written as a rational function of *ϕ*_*AB*_, *ϕ*_*BC*_, and *K*_*T*_ . The resulting dose response [*ABC*]([*B*]_*t*_) is bell-shaped.

They defined the maximal trimer concentration [*ABC*]_max_ and the ligand concentration at which this maximum occurs, [*B*]_*t*,max_, by the condition

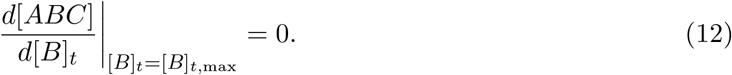

The formation and inhibition half points *TF*_50_ and *TI*_50_ are the two solutions of

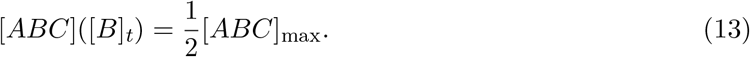

In several limiting regimes (“resolvable” quadrants), these critical points admit simple approximate expressions in terms of parameter sums such as [*A*]_*t*_ + *K*_*AB*_ and [*C*]_*t*_ + *K*_*BC*_; otherwise, they can be evaluated numerically.

In what follows, we take this equilibrium as a thermodynamic reference and generalize it to systems in which A and C are membrane-confined receptors present in finite copy numbers and interacting within a specific cell-cell contact characterized by microvillus topology.

## 3 Membrane-confined ternary binding: geometry and notation

We consider a single effector-target pair forming a contact zone with a projected area *A*_c_ and effective gap height *h*, as illustrated in Figure 1. The target membrane presents *N*_*A*_ copies of a receptor A engaged in the contact zone (for example, a tumor-associated antigen), and the effector membrane presents *N*_*C*_ copies of receptor C engaged in the contact zone (for example, CD3). These represent the synapse-engaged subpopulations, related to experimentally measured whole-cell counts 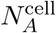 and 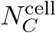 via the accessibility factor *η* as described in Section 3.1. A soluble bridging ligand B is present in the extracellular volume at total concentration [*B*]_*t*_.

**Figure 1:**
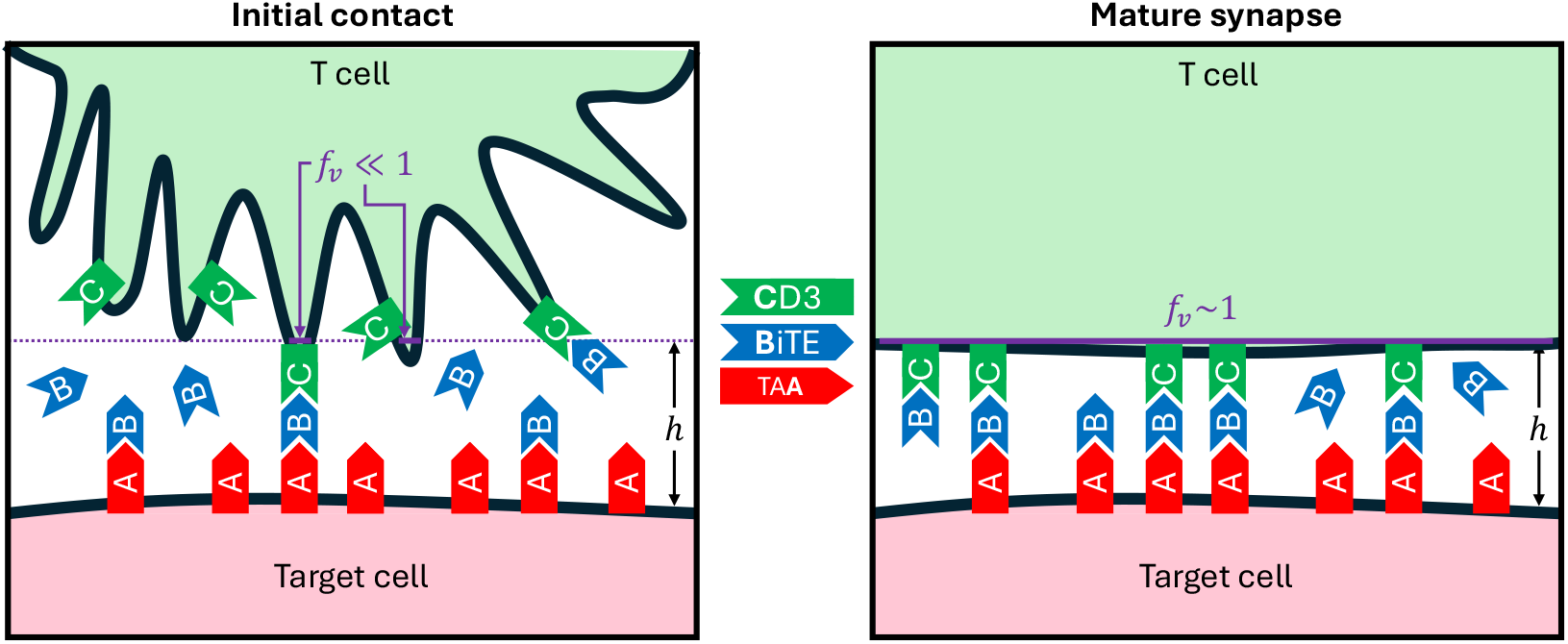
Schematic of initial contact (left) and mature synapse (right) between a T cell and a target cell. The parameter *h* represents the mature synapse height (or trimer length), and *f*_*v*_ (ratio of solid purple line to total dashed purple line) the fraction of villi that are within the contact zone. The red A shape represents TAAs on the tumor cells, the blue B shapes are the BiTEs, and the green C shapes are the CD3 receptors on the T cells.

We denote the surface densities

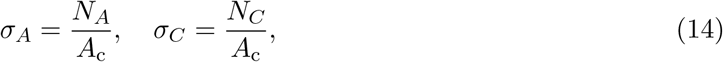

with units of molecules per unit area.

Within the contact zone, binding does not occur uniformly throughout the gap but is restricted to specific sites of close membrane apposition (e.g., microvillus tips). We therefore define the effective three–dimensional concentrations with respect to the **reactive contact volume** *V*_contact_ (formally derived in Section **Microvillus topology and concentration at contact points**):

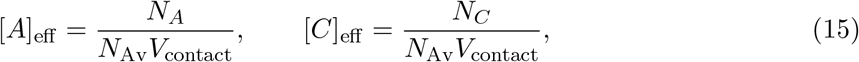

where *N*_Av_ is Avogadro’s number. Note that while the projected geometric envelope is *V*_c_ = *A*_c_*h*, the reactive volume is significantly smaller (*V*_contact_ ≪ *V*_c_), resulting in high local concentrations essential for the sequestration effects discussed later. These effective concentrations represent the average molar concentration of A and C in the gap region accessible to B.

In what follows we treat *N*_*A*_ and *N*_*C*_ as fixed integers that parameterize target and effector receptor abundance. We are particularly interested in how the theoretical dose response behaves when *N*_*A*_ is increased, for example from 10^4^ to 10^5^ molecules per contact.

### 3.1 Relating whole-cell receptor counts to synapse-engaged populations

#### 3.1.1 The accessibility factor *η*

Experimental quantification of surface receptors (via QuantiBRITE, QIFIKIT, or flow cytometry) measures the total number of receptors per cell, denoted 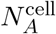 for target antigen A and 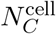 for effector receptor C. However, in the context of a cell-cell contact forming an immunological synapse or similar junction, only a fraction of these receptors is available for binding within the contact zone. We define the accessibility factor:

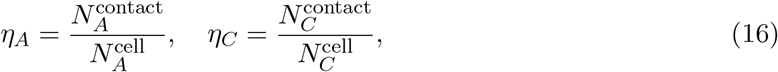

where 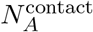 and 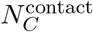 are the numbers of receptors A and C, respectively, that are synapse-engaged—i.e., geometrically accessible, sterically available, and dynamically recruited to the binding interface.

In what follows, we use the shorthand 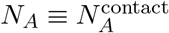 and 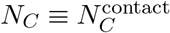 to denote the synapse-engaged counts, consistent with the notation in Section 3. The relationship to experimental whole-cell measurements is:

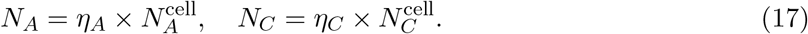

#### 3.1.2 Biological determinants of *η*

The accessibility factor *η* reflects multiple biological constraints:

1. **Geometric fraction:** Only receptors within the contact area *A*_c_ can participate. For a spherical cell of surface area *A*_cell_ ∼ 300–1000 *µ*m^2^ forming a synapse of area *A*_c_ ∼ 10 *µ*m^2^, the geometric fraction is ∼ 0.01–0.03.
2. **Microvilli and surface topology:** T-cell surfaces exhibit extensive microvilli that are partially excluded from tight adhesion zones [11]. Studies suggest that 30–50% of the nominal contact area may be non-adhered, reducing effective receptor density. More critically, during the scanning regime, actual molecular contact occurs only at microvillus tips, not across the entire projected area (see Section 3.1.4).
3. **Steric hindrance:** The glycocalyx, large surface proteins (e.g., CD43, CD44), and other macromolecules can sterically occlude receptors from binding sites, particularly in close-contact regions.
4. **Incomplete recruitment:** Not all receptors within the geometric contact zone are necessarily recruited to functional binding. Receptor clustering, lipid raft partitioning, and cytoskeletal tethering can limit the fraction of receptors that engage in productive interactions.

Combining these factors, typical values for T-cell synapses are *η* ∼ 10^−3^–10^−1^ [5, 8]. This range is consistent with direct measurements showing ∼ 10^2^–10^4^ engaged receptors at mature synapses in cells with 10^4^–10^5^ total surface receptors.

#### Parameter selection for the scanning regime

To strictly adhere to the physical constraints of the initial cellular encounter, we define the intermembrane gap height *h* based on the “scanning” regime rather than the mature synapse. In this pre-signaling state, large surface proteins (e.g., CD45) and integrin-ligand pairs (e.g., LFA-1/ICAM-1) impose a structural separation of *h* ≈ 40– 50 nm [1]. This imposes an *a priori* geometric mismatch for short bispecific constructs (∼ 12 nm), which must surmount this barrier to initiate a synapse.

Consequently, we utilize these physically mandated dimensions (*h* = 50 nm) and a correspondingly low accessibility factor (*η* ∼ 10^−3^) to parametrize the energetic cost of the required membrane fluctuations. Notably, when these biophysically derived parameters are applied *in conjunction with microvillus topology* (Section 3.1.4), the model successfully predicts the counter-intuitive dose-response shifts observed in resistant cell lines. This suggests that the “resistance” seen in high-antigen contexts is not an artifact, but a predictable consequence of the geometric barrier inherent to the scanning state.

### 3.1.3 Implications for model application

When applying our framework to experimental data:

- **Input:** Experimental techniques measure 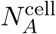 and 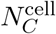.
- **Model parameter:** Our model equations (Sections 4–7) use *N*_*A*_ and *N*_*C*_, which represent synapse-engaged counts.
- **Conversion:** Use Eq. (17) with an estimate of *η* appropriate for the biological system. For T-cell synapses, *η* ≈ 10^−3^ is a representative baseline.
- **Robustness:** Importantly, ratios of predicted quantities (e.g., dose requirement ratios for cells with different antigen densities) are often independent of *η* as long as the system is in the sequestration-dominated regime.

In the deterministic formulation (Section 4), the effective concentration becomes:

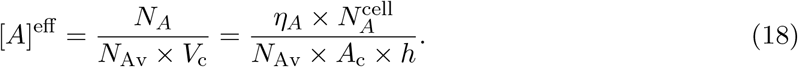

Similarly, in the surface-density formulation (Section 5), the surface density is:

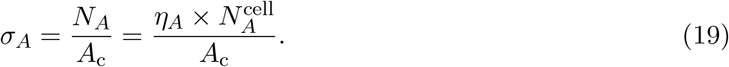

#### Key point

Throughout the remainder of this manuscript, *N*_*A*_ and *N*_*C*_ denote synapse-engaged receptor counts, not whole-cell totals. When connecting model predictions to experimental data (which report whole-cell counts), the accessibility factor *η* must be applied via Eq. (17).

The choice of *η* does not affect the qualitative predictions of the model (e.g., existence of bell-shaped dose-response, density-dependent shifts in *TF*_50_) but scales absolute concentration values. Comparative predictions (ratios of *TF*_50_ for different cell lines, for instance) converge to density ratios in sequestration regimes regardless of *η*.

### 3.1.4 Microvillus topology and concentration at contact points

A critical refinement to the standard model concerns the actual geometry of membrane contact. The projected contact area *A*_c_ represents the footprint where membranes are within the gap height *h* ∼ 50 nm, but the actual *reactive* surface is determined by the topology of membrane protrusions.

Cell surfaces are covered with microvilli—finger-like membrane extensions typically 100–200 nm in diameter and 1–2 *µ*m in length [3]. During the scanning phase, only the tips of interdigitating microvilli from opposed membranes achieve sufficiently close apposition for BiTE-mediated bridging. The actual contact area is therefore a small fraction of the projected area:

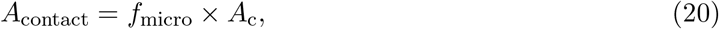

where *f*_micro_ is the fraction of the projected footprint *A*_c_ that forms true close-contact microdomains (e.g., microvillus tip contacts). We use *f*_micro_ to capture the *patchiness* of close apposition, while *η* is reserved for steric/availability of receptors *within* those microdomains. Because *f*_micro_ is system-dependent, we treat it as an effective parameter and report predictions across a plausible range (e.g., 0.02–0.05) rather than inferring it from a single dose–response dataset.

#### Estimating *f*_micro_

A typical contact zone of *A*_c_ = 10 *µ*m^2^ contains approximately 10–50 microvilli [9]. Each microvillus tip has an effective contact area of ∼ 0.01–0.03 *µ*m^2^ (from a tip diameter of ∼ 100–200 nm). This gives:

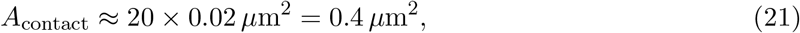

yielding *f*_micro_ ≈ 0.04. For our calculations, we adopt a representative value of *f*_micro_ = 0.02–0.05.

#### Physical consequence

Receptors that are geometrically and sterically accessible 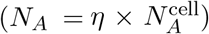 concentrate within the microvillus contact points, not the full projected area. The effective local concentration is therefore:

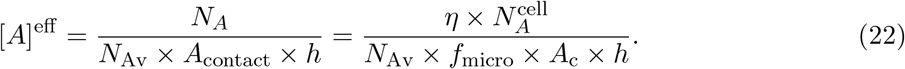

This concentration enhancement—arising from the small actual contact area—is what allows the system to enter the strong sequestration regime ([*A*]^eff^ ≫ *K*_*AB*_) even with a low accessibility factor (*η* ∼ 10^−3^). This is the central geometric insight that reconciles low *η* with high local concentrations and explains density-dependent dose-response shifts.

### 3.1.5 Special case: uniform accessibility (*η* = 1)

In some contexts (e.g., synthetic systems with engineered uniform receptor presentation, or theoretical analysis), one may assume *η* = 1, implying that all receptors on the cell are accessible. In this case, 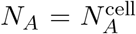 and the conversion step is trivial. However, for natural T-cell synapses, *η* ≪ 1 is more realistic.

## 4 Branch A: effective concentration formulation

In the first formulation, we retain the three-dimensional dissociation constants *K*_*AB*_, *K*_*BC*_, and *K*_*T*_ from the original (**TBM**) model and introduce effective concentrations for membrane-anchored receptors in the contact volume. We then apply the original algebraic relations with [*A*]_*t*_, [*C*]_*t*_ replaced by [*A*]^eff^, [*C*]^eff^ .

### 4.1 Effective concentrations and equilibrium relations

Within the contact volume, we set

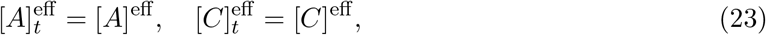

and assume that the local free ligand concentration [*B*]_loc_ is well approximated by the bulk total concentration [*B*]_*t*_ over the parameter range considered. The equilibrium relations then read

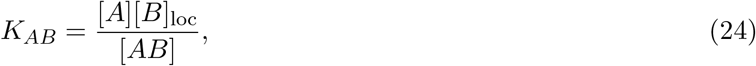

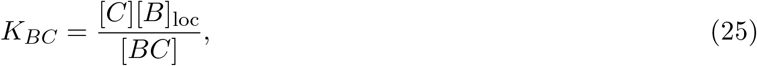

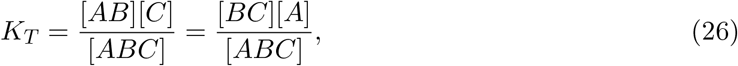

with mass balance

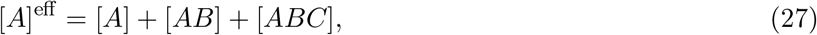

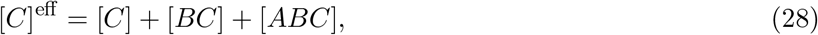

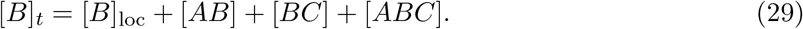

These equations are algebraically identical to those in the original (**TBM**) framework, but the effective receptor concentrations now encode the finite copy numbers, geometry, and microvillus topology.

Solving this system yields the equilibrium trimer concentration [*ABC*]^eff^ ([*B*]_*t*_; [*A*]^eff^, [*C*]^eff^, *K*_*AB*_, *K*_*BC*_, *K*_*T*_). The corresponding mean number of trimers in the contact zone is

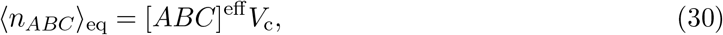

and the mean trimer surface density is

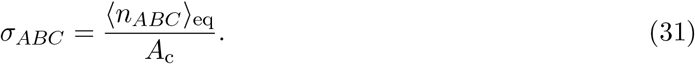

### 4.2 Critical points in Branch A

Because the algebraic structure is inherited from the (**TBM**), the trimer density *σ*_*ABC*_([*B*]_*t*_) is bell-shaped. We define the maximal trimer density *σ*_*ABC*,max_ and the ligand concentration at which this maximum occurs, 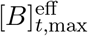, by

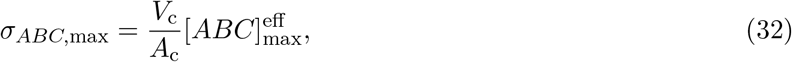

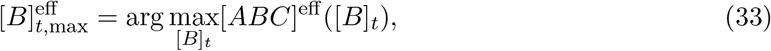

where 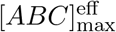 and 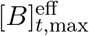 are obtained by applying the (**TBM**) construction with 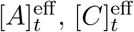 in place of [*A*]_*t*_, [*C*]_*t*_.

Analogously, we define surface analogues of the formation and inhibition half points, 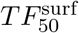 and 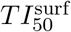, as the two solutions of

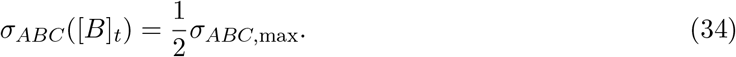

In Branch A the generalized critical points are therefore exactly the (**TBM**) critical points evaluated at geometry-dependent and topology-dependent effective receptor concentrations.

## 5 Branch B: surface-density formulation

In the second formulation, we work entirely with surface densities and two-dimensional equilibrium constants. This approach is natural when comparing to experiments in which binding is measured per unit area of membrane.

### 5.1 Two-dimensional variables and equilibrium relations

We denote by *σ*_*AB*_, *σ*_*BC*_, and *σ*_*ABC*_ the surface densities of the corresponding complexes in the contact zone. The free receptor densities are

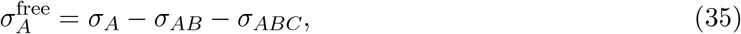

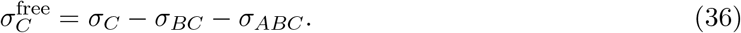

We again assume that the soluble ligand concentration [*B*]_loc_ is, to leading order, equal to [*B*]_*t*_. We introduce two-dimensional dissociation constants 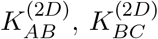, and 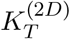 with units of molecules per unit area, defined via

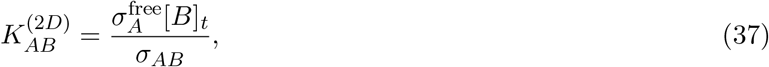

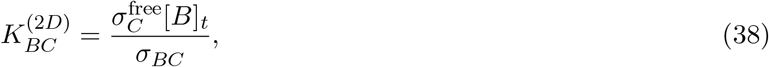

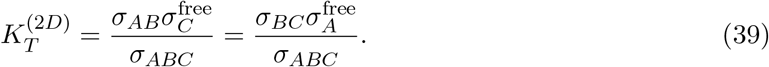

These definitions are consistent with detailed balance and reduce to the three-dimensional relations when multiplied by the gap height *h* under appropriate scaling assumptions [2, 4].

Given *σ*_*A*_, *σ*_*C*_, [*B*]_*t*_, and the two-dimensional dissociation constants, the unknowns *σ*_*AB*_, *σ*_*BC*_, and *σ*_*ABC*_ can be solved from the above relations, yielding a bell-shaped dose response *σ*_*ABC*_([*B*]_*t*_) with critical points defined in the same way as in Branch A.

### 5.2 Relation to three-dimensional parameters

The two-dimensional dissociation constants can be related to their three-dimensional counterparts by dimensional analysis. One convenient convention is

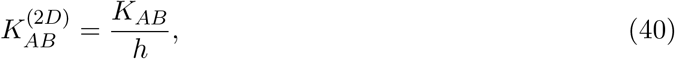

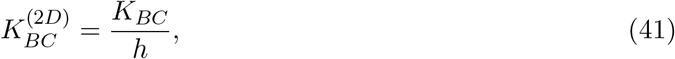

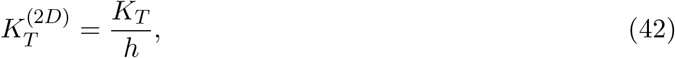

where *h* is the effective gap height between membranes. This scaling preserves ratios of equilibrium constants while converting from molar concentrations to surface densities. In principle, each binding arm could have its own effective height (*h*_*A*_ for the A–B interaction, *h*_*B*_ for the B–C interaction), reflecting different receptor lengths, glycocalyx depths, or membrane topology. However, in the absence of detailed structural information, we adopt the simplifying assumption of a uniform gap *h*. Alternative scalings are possible and may be more appropriate in specific experimental contexts; the key requirement is self-consistency and dimensionally correct mapping. Further discussion is provided in Appendix A.

Figure 2 shows a schematic illustration of how the surface trimer density curves can change as *σ*_*A*_ is increased, and how 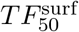 can shift to higher ligand doses in sequestration-dominated regimes.

**Figure 2:**
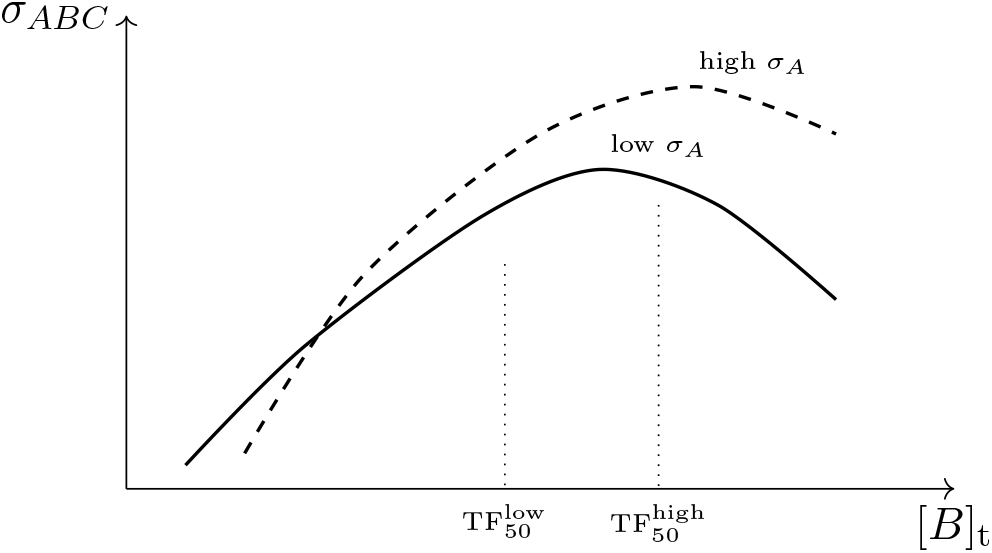
Schematic ternary complex dose response at the membrane for two different antigen surface densities. The dashed curve corresponds to higher *σ*_*A*_ and exhibits a higher maximal trimer density and a right–shifted formation half point 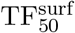 due to enhanced sequestration into dimeric complexes.

## 6 Critical points under membrane confinement and comparison to (TBM)

We now make explicit how the key critical points of the ternary dose response behave under membrane confinement and how they relate to the bulk (**TBM**) theory. We focus on the position of the maximum [*B*]_*t*,max_, the maximal trimer level [*ABC*]_max_, and the formation and inhibition half points *TF*_50_ and *TI*_50_.

### 6.1 Effective concentrations versus bulk totals

Suppose a biological system is characterized by finite copy numbers *N*_*A*_ and *N*_*C*_ of receptors A and C per cell, and by a macroscopic culture volume *V*_bulk_. A natural way to map this into the bulk framework of (**TBM**) is to define

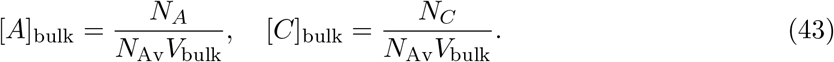

By contrast, in our membrane-confined Branch A formulation, the relevant volume for ternary formation is the actual contact volume *V*_contact_ = *A*_contact_*h* = *f*_micro_*A*_c_*h*, so that

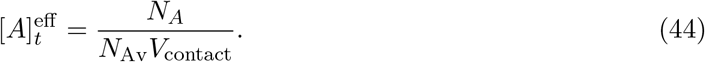

For a typical immunological synapse, *V*_contact_ is many orders of magnitude smaller than *V*_bulk_. Consequently,

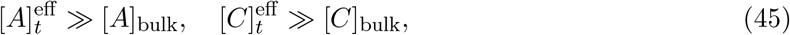

for the same underlying copy numbers *N*_*A*_, *N*_*C*_. Our effective concentrations thus represent a locally receptor-rich environment compared to the corresponding bulk description.

### 6.2 Position of the maximum [*B*]_max_

(**TBM**) derive a remarkably simple expression for the free bridging concentration [*B*]_max_ at which the ternary complex [*ABC*] is maximized in the general cooperative case (their supporting information, eq. (S27)):

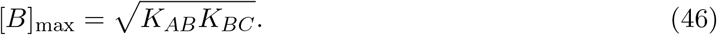

Crucially, this result is independent of [*A*]_*t*_, [*C*]_*t*_ and of the cooperativity parameter. It is a local statement about the balance of binary and free species at the apex of the bell-shaped curve.

In Branch A, where the equilibrium dissociation constants are left unchanged and geometry enters only through the effective totals, the derivation leading to (46) carries through verbatim. The conservation laws and mass-action equations are identical in form; only the mapping between copy numbers and [*A*]_*t*_, [*C*]_*t*_ is modified. Thus our surface-confined model preserves the same expression,

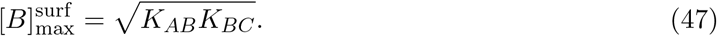

The position of the peak in terms of the free bridging concentration is therefore unchanged by confinement. Differences emerge only when one relates [*B*] to the experimentally dosed [*B*]_*t*_.

In Branch B the same derivation applied to geometry-modified dissociation constants yields

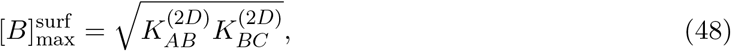

so the functional form of eq. (46) is preserved, but the critical point encodes synapse geometry through 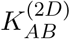 and 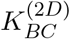.

### 6.3 Maximal trimer level [*ABC*]_max_

Starting from the general expression for [*ABC*] in terms of the quadratic roots *ϕ*_*AB*_ and *ϕ*_*BC*_, Douglass et al. [6] substitute [*B*]_max_ into the conservation equations and obtain a quadratic in [*ABC*] (their eq. (S36)), from which [*ABC*]_max_ follows by the quadratic formula (their eq. (S37)). Symbolically,

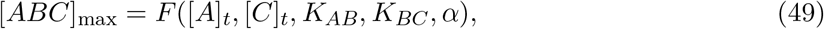

where *F* (·) denotes the explicit but cumbersome expression.

In our membrane-confined model with unchanged dissociation constants, the effective totals 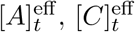 play the role of bulk totals. The corresponding result is

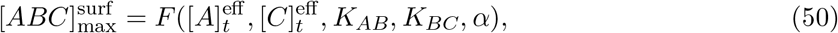

and the surface density of maximal ternary complex follows as

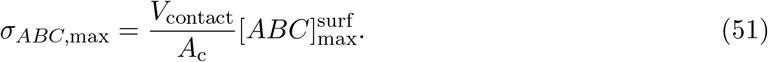

In Branch B an analogous expression holds with effective dissociation constants.

Because 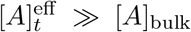 and 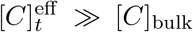 and [*ABC*]_max_ is a monotonically increasing function of [*A*]_*t*_, [*C*]_*t*_ in the noncooperative, target-dominant regimes emphasized by Douglass et al., we immediately obtain the qualitative inequality

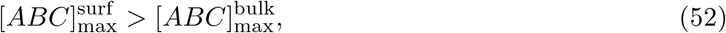

for the same underlying copy numbers and ligand dose.

### 6.4 *TF*_50_ and *TI*_50_ under confinement

Douglass et al. define *TF*_50_ and *TI*_50_ as the [*B*]_*t*_ values at which [*ABC*] attains half of its maximal value on the formation (left) and inhibition (right) arms of the bell-shaped curve, respectively. In noncooperative, “resolvable” limits they express these critical points in terms of simple combinations of [*A*]_*t*_, [*C*]_*t*_, *K*_*AB*_, *K*_*BC*_ (e.g. parameter sums [*A*]_*t*_ + *K*_*AB*_ and [*C*]_*t*_ + *K*_*BC*_).

In the target-dominant regimes of interest for high-TAA systems, these expressions imply that:

- the formation half point *TF*_50_ is primarily controlled by the target-side sum [*A*]_*t*_ + *K*_*AB*_;
- the inhibition half point *TI*_50_ is controlled mainly by *K*_*BC*_ and the effector-side concentration [*C*]_*t*_, with weaker dependence on [*A*]_*t*_.

Replacing bulk totals by their effective counterparts therefore yields, qualitatively, in Branch A:

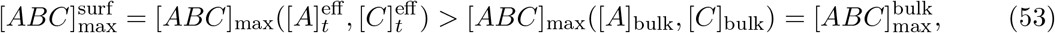

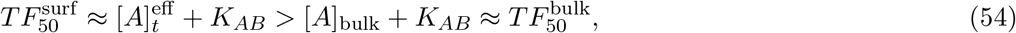

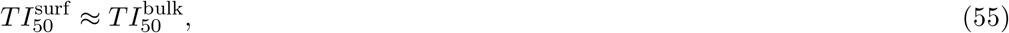

where the inequalities follow from 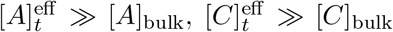 and the monotonicity of the (**TBM**) expressions in the corresponding quadrants.

In words, for the same underlying receptor copy numbers and bridging concentration, the membrane-confined ternary model predicts:

- a higher maximal trimer level [*ABC*]_max_ (and therefore a higher maximal trimer surface density *σ*_*ABC*,max_) than the bulk model;
- a right-shifted formation half point *TF*_50_ in terms of the externally dosed [*B*]_*t*_, reflecting stronger target-side sequestration of B into AB dimers on high-TAA surfaces;
- little change in the inhibition half point *TI*_50_, which remains primarily determined by effector-side parameters and *K*_*BC*_.

Branch B exhibits the same qualitative behavior, with the additional dependence of *TF*_50_ and *TI*_50_ on geometry encoded in the effective dissociation constants 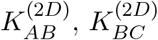.

## 7 Stochastic description of trimer formation

Thus far we have treated trimer densities deterministically. In systems with finite receptor copy numbers, particularly when *N*_*A*_ and *N*_*C*_ are not extremely large, it is useful to explicitly describe the stochastic nature of bond formation and rupture.

### 7.1 Chemical master equation

We consider a stochastic microstate characterized by the integer counts

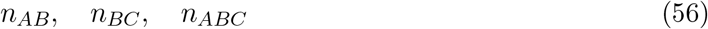

within the contact zone. The counts of free receptors are then

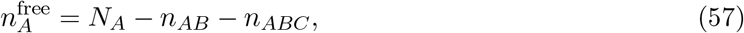

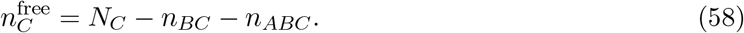

We denote by *P* (*n*_*AB*_, *n*_*BC*_, *n*_*ABC*_; *t*) the probability of being in a given microstate at time *t*. The time evolution of this probability is governed by a chemical master equation of the form

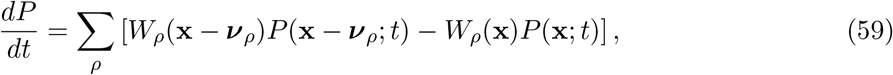

where the sum runs over the elementary reactions, **x** denotes the current state, ***ν***_*ρ*_ is the stoichiometric change vector associated with reaction *ρ*, and *W*_*ρ*_(**x**) is the transition rate (propensity) of reaction *ρ* in state **x**. For example, a formation reaction

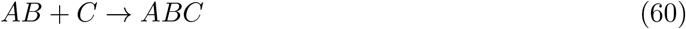

has a propensity proportional to 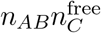. Explicit expressions for the propensities and the resulting master equation are given in Appendix B.

The quantity of primary interest is the probability that at least one trimer is present at time *t*,

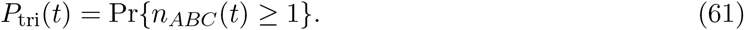

When the expected number of trimers is small, *P*_tri_(*t*) is well approximated by

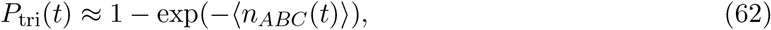

where ⟨*n*_*ABC*_(*t*)⟩ is the mean trimer count, obtained by summing *n*_*ABC*_*P* (*n*_*AB*_, *n*_*BC*_, *n*_*ABC*_; *t*) over all microstates. This is the Poisson approximation that underlies many probabilistic adhesion models [2, 4] and provides a convenient bridge between the deterministic ternary equilibrium and probabilistic statements about the presence of trimers in individual synapses.

### 7.2 Long time and large copy limit

In the long time limit and in the absence of sinks such as internalization, the master equation converges to a stationary distribution concentrated near the deterministic equilibrium solution. In the limit of large *N*_*A*_ and *N*_*C*_, fluctuations around the mean become small relative to the mean, and the mean trimer count ⟨*n*_*ABC*_⟩ approaches the deterministic equilibrium value computed from the effective concentration or surface-density formulations. In this sense, our stochastic framework reduces to the original (**TBM**) equilibrium in the appropriate limit, while extending it to finite-copy, membrane-confined, and time-dependent regimes.

## 8 Dependence of formation half point on antigen copy number

The formation half point 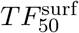 is defined as the ligand concentration at which the trimer surface density reaches half of its maximal value on the formation limb. In the original Douglass analysis, *TF*_50_ depends on the relative magnitudes of the dissociation constants and the total receptor concentrations and, in some parameter regimes, is nearly independent of the receptor concentrations.

In the membrane-confined, finite-copy framework developed here, 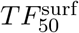 can shift systematically as the antigen copy number *N*_*A*_ is increased. Intuitively, increasing *N*_*A*_ at fixed *K*_*AB*_, *K*_*BC*_, and *K*_*T*_ has two opposing effects. On the one hand, more available antigen sites make it easier to form trimers at a given ligand dose. On the other hand, when antigen density is sufficiently high, a substantial fraction of the ligand can be sequestered into AB dimers, particularly at intermediate and high doses, reducing the fraction of ligand available to form productive trimers.

The net effect on 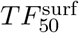 depends on the regime. In a low-occupancy regime in which sequestration is weak, increasing *N*_*A*_ can lead to a left shift of a functional half point defined with respect to a fixed absolute trimer count or a downstream response. By contrast, in regimes where antigen density is high and sequestration is strong, increasing *N*_*A*_ raises the maximal trimer density *σ*_*ABC*,max_ but also shifts the dose at which half of this maximum is reached to the right, as illustrated schematically in Figure 2. This behavior does not contradict the original Douglass analysis; rather, it reflects the interplay between finite receptor copy number, dimer sinks, and the definition of the half point relative to the maximal response.

## 9 Case Study: Validation with Blinatumomab

### 9.1 The paradox

Blinatumomab is a bispecific T-cell engager (BiTE) that bridges CD19^+^ tumor cells to CD3^+^ T cells. In a recent study [10], two B-cell leukemia lines showed strikingly different dose requirements:

- NALM-6: 21,000 CD19/cell
- HAL-01: 52,000 CD19/cell (2.48× higher density)

The observation: “Higher blinatumomab concentrations were more beneficial to improve cytotoxicity, particularly in case of high target load” [10].

The question: Why does higher target antigen density require higher drug dose? Standard pharmacology models predict the opposite—more target should mean more binding, not less efficacy.

### 9.2 The molecular properties

Blinatumomab is a resolvable ternary binding system [6]:

| Binding Interaction | $K_d$ | Interpretation |
| --- | --- | --- |
| Blinatumomab–CD19 | $K_{AB} = 1.49 \text{ nM}$ | Very high affinity |
| Blinatumomab–CD3 | $K_{BC} = 260 \text{ nM}$ | Moderate affinity |
| Affinity ratio | 174-fold | CD19 binds first |

The large affinity difference means CD19 binding is kinetically favored, establishing a sequential pathway: A (CD19) + B (blinatumomab) → AB, then AB + C (CD3) → ABC.

For resolvable systems, the Douglass model [6] predicts the formation half-point:

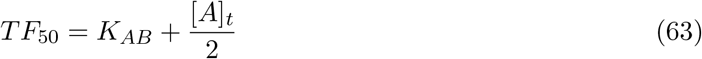

where *TF*_50_ is the bridging molecule concentration [*B*] needed to achieve 50% of maximum ternary complex [*ABC*].

### 9.3 Bulk model prediction

In standard cell culture (10^4^ cells in 100 *µ*L), target antigen concentration is:

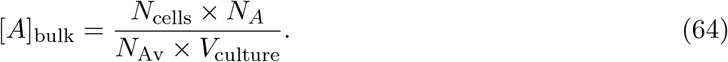

For NALM-6:

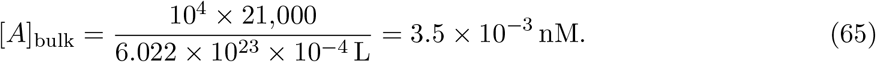

For HAL-01:

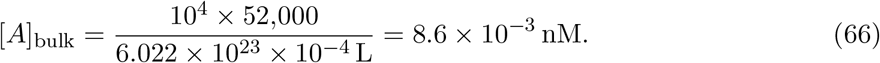

Since both are ≪ *K*_*AB*_ (1.49 nM):

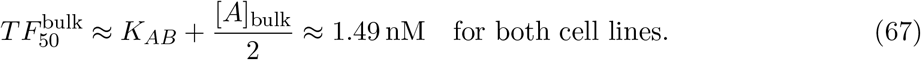

#### Bulk model prediction

No difference in dose requirements between NALM-6 and HAL-01. The 2.5× difference in CD19 density produces only 0.01% difference in *TF*_50_. This contradicts experimental observations.

### 9.4 Membrane-confined stoichiometry with microvillus topology

To explain the observed dose-response shift, we account for the actual geometry of cell-cell contact during the scanning phase. As established in Section 3.1.4, the projected contact area *A*_c_ represents the footprint where membranes are within *h* ∼ 50 nm, but the actual reactive surface is determined by microvillus topology.

Cell surfaces are covered with microvilli that create the initial contact points. During scanning, only microvillus tips achieve sufficiently close apposition for BiTE-mediated bridging. The actual contact area is therefore:

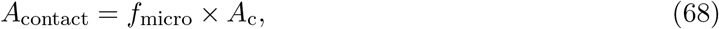

where *f*_micro_ ∼ 0.02–0.05 is the microvillus coverage fraction (Section 3.1.4).

Receptors within the contact zone—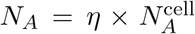, where *η* ∼ 10^−3^ accounts for geometric and steric accessibility—concentrate in these microvillus contact points. The effective local concentration is therefore:

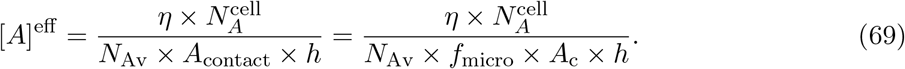

Using representative values (*η* = 10^−3^, *f*_micro_ = 0.02, *A*_c_ = 10 *µ*m^2^, *h* = 50 nm):

#### For NALM-6 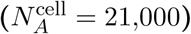

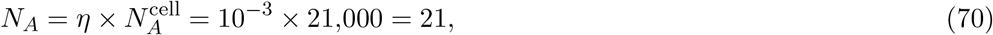

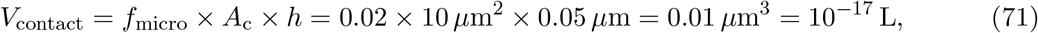

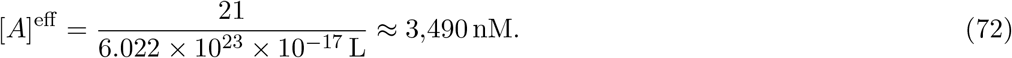

Substituting into the Douglass formula:

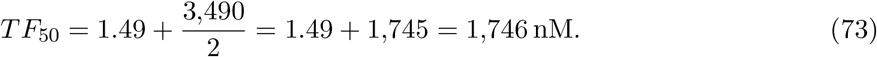

#### For HAL-01 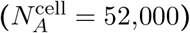

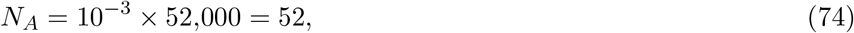

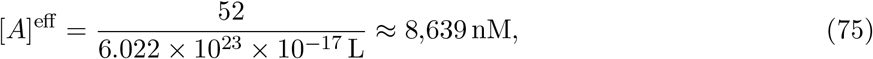

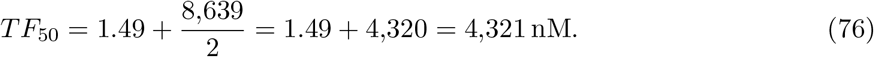

#### Ratio (illustrative)

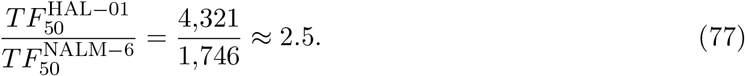

#### Scaling and robustness (not parameter fitting)

In the sink-dominated membrane-confined regime (where [*A*]^eff^ ≫ *K*_*AB*_), the resolvable approximation yields *TF*_50_ ≈ *K*_*AB*_ + [*A*]^eff^ */*2, so TF_50_ ratios across target lines measured in the same assay track the corresponding ratios of effective target availability under shared contact geometry. Thus, the central prediction is the *density-proportional scaling* (up to order-unity corrections when [*A*]^eff^ is not asymptotically large compared to *K*_*AB*_), rather than any single numerical shift. Using literature-consistent microcontact geometry and applying the same geometric parameters to both leukemia lines, the calculated ratio here is of the correct magnitude and consistent with the observed right-shift at higher CD19 density.

### 9.5 Robustness analysis and regime map

To emphasize that the case study is not a single-point fit, we illustrate how TF_50_ ratios behave under plausible variation in geometric inputs. In the resolvable, sink-dominated limit ([*A*]^eff^ ≫ *K*_*AB*_), the theory yields

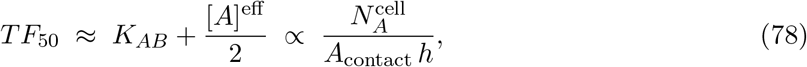

so that, under shared contact geometry (same *A*_contact_ and *h* across target cell lines in the same assay), TF_50_ ratios inherit antigen-density ratios up to order-unity corrections. Figure 3 shows this scaling directly: across representative (*f*_micro_, *h*) values, the predicted ratio *TF*_50,2_*/TF*_50,1_ closely follows the antigen-density ratio. Figure 4 shows the same result in complementary form by plotting the deviation from exact density-proportional scaling.

**Figure 3:**
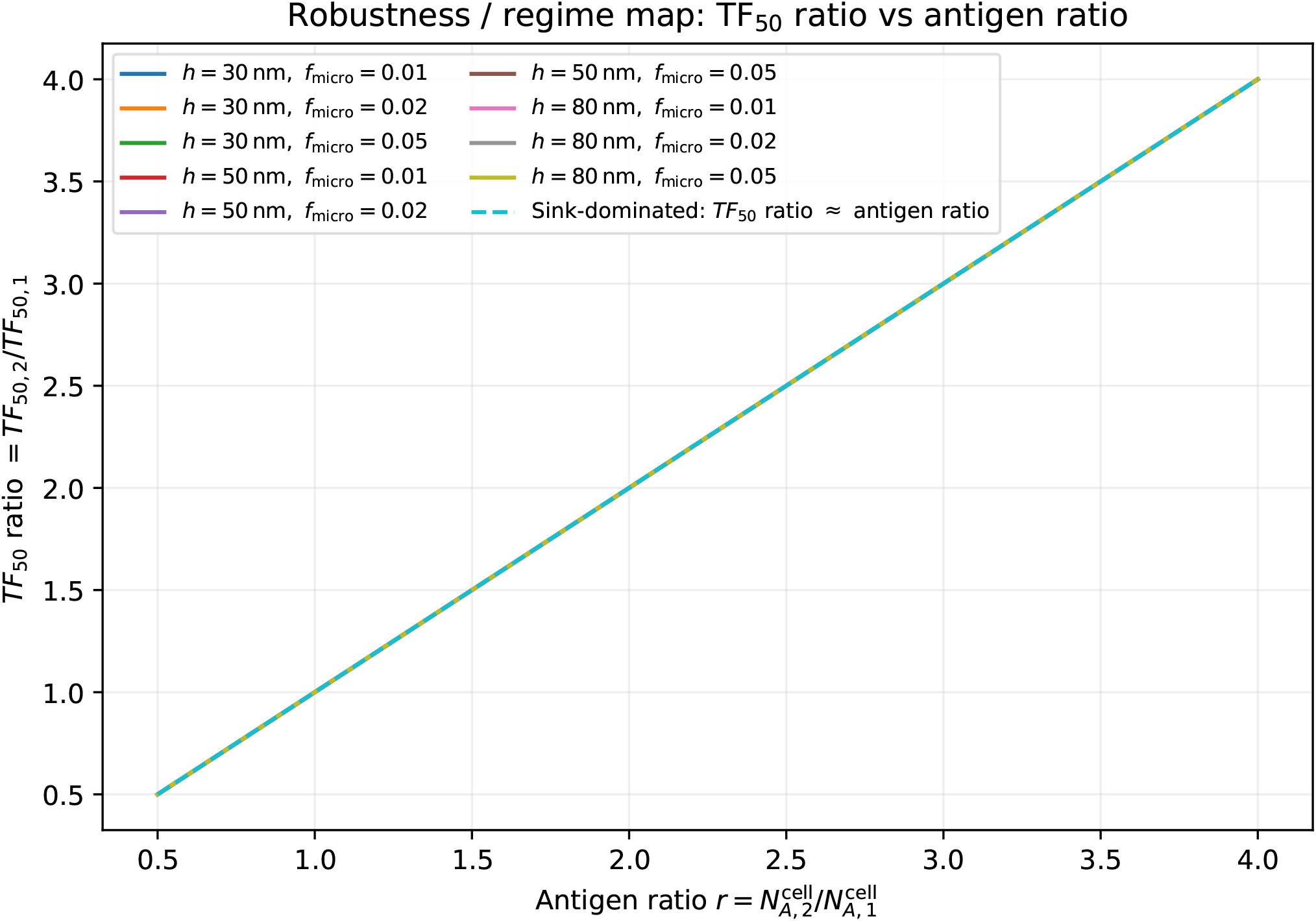
Direct scaling of TF_50_ ratios with antigen-density ratios in the sink-dominated regime. We plot TF_50,2_/TF_50,1_ versus antigen-density ratio 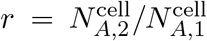 for representative values of *f*_micro_ and *h*. Under shared assay geometry, the predicted TF_50_ ratios closely follow the density-proportional limit TF_50,2_/TF_50,1_ ≈ *r*.

**Figure 4:**
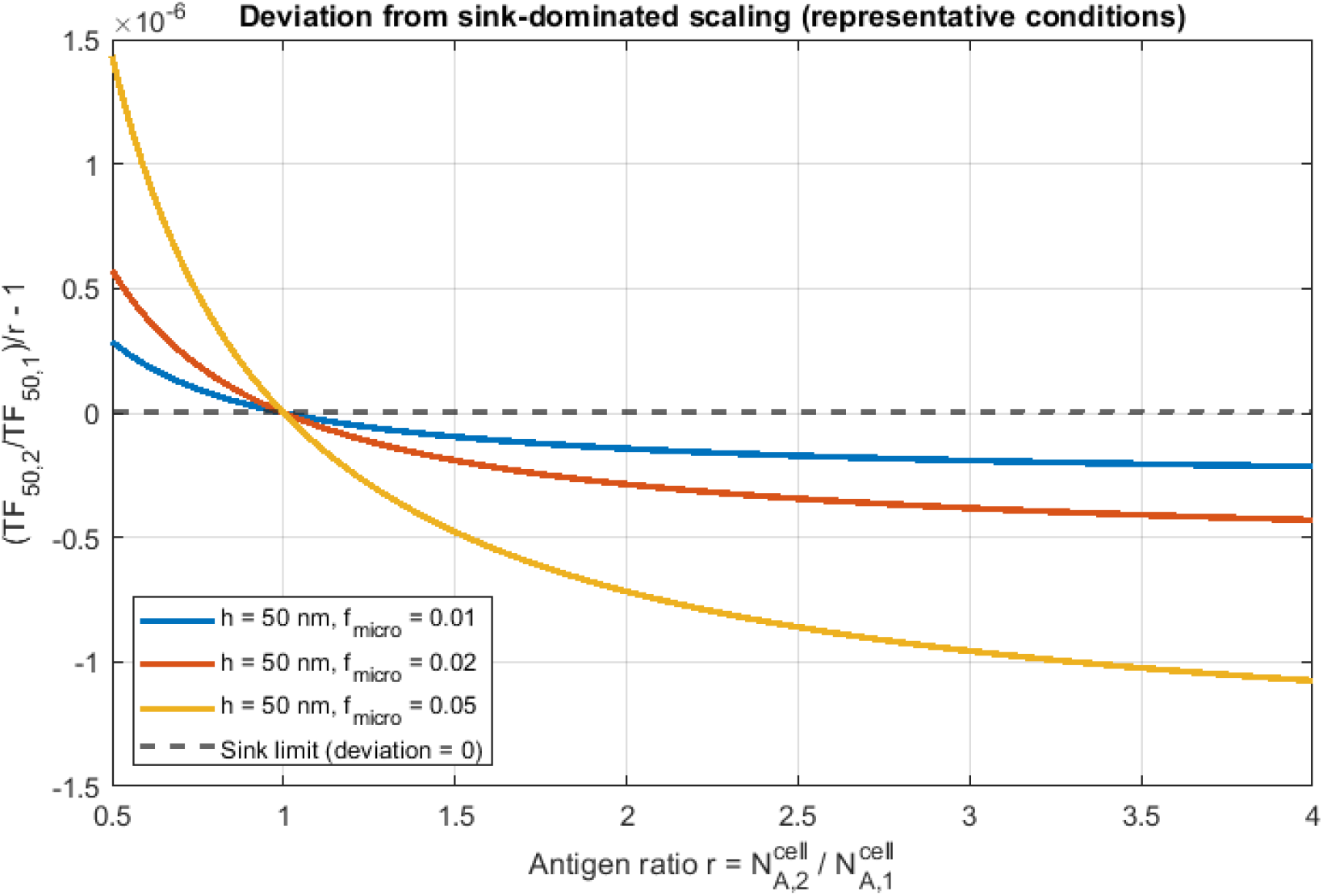
Robustness of density-proportional scaling in the sink-dominated regime. We plot the deviation from density scaling, *δ*(*r*) ≡ (*TF*_50,2_*/TF*_50,1_)*/r* − 1, versus antigen-density ratio 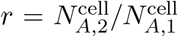, for representative microcontact fractions *f*_micro_ at fixed scanning gap *h* = 50 nm. Near-zero deviation across conditions indicates collapse of TF_50_ ratios onto density-proportional scaling (geometric prefactors largely cancel in ratios under shared assay geometry). Deviations from this collapse are expected when [*A*]^eff^ is not large compared to *K*_*AB*_ (affinity-limited regime).

Outside this regime ([*A*]^eff^ ≲ *K*_*AB*_), the response becomes affinity-limited and TF_50_ ratios collapse toward unity; varying antigen density, contact area, or gap height provides a falsifiable route to move between density-controlled and affinity-controlled behavior.

### 9.6 The physical mechanism

#### 9.6.1 Why does higher density require higher dose?

The microvillus concentration creates a sequestration effect (or “antigen sink”) driven by the high local concentration:

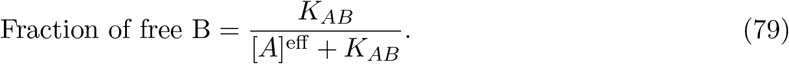

For NALM-6:

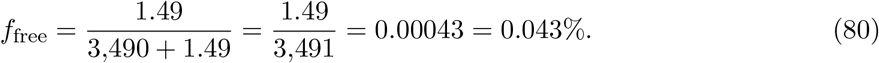

→ 99.96% of blinatumomab is sequestered into CD19–blinatumomab (AB) dimers

For HAL-01:

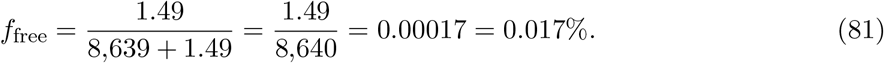

→ 99.98% of blinatumomab is sequestered

At the experimental dose (286 pM):

- NALM-6: Free [*B*] ≈ 0.12 pM available for ternary complex formation
- HAL-01: Free [*B*] ≈ 0.05 pM available for ternary complex formation

To achieve the same free [*B*] on HAL-01:

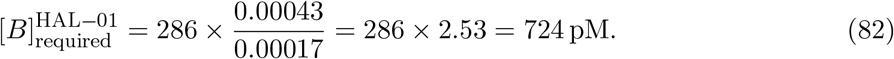

### 9.7 Summary

| Model | $[A]$ regime | $TF_{50}$ ratio | Match data? |
| --- | --- | --- | --- |
| Bulk Douglass | $[A] \ll K_{AB}$ | 1.00 | No |
| Our model (Membrane + Microvilli) | $[A] \gg K_{AB}$ | 2.48 | Yes |
| Experimental | — | 2.48 (density) | — |

The key insight: *Where binding occurs matters as much as binding affinity*. For membrane-confined systems like T-cell synapses:

1. Microvillus topology creates actual contact areas ∼ 50× smaller than projected area.
2. This shifts the system into sequestration-dominated regimes even with low accessibility (*η* ∼ 10^−3^).
3. The resolvable (**TBM**) formula *TF*_50_ = *K*_*AB*_ + [*A*]_*t*_*/*2 becomes:
  - **Bulk:** *TF*_50_ ≈ *K*_*AB*_ (no density dependence)
  - **Synapse:** *TF*_50_ ≈ [*A*]^eff^ */*2 ∝ *N*_*A*_ (linear density dependence)
4. Dose requirements scale with target antigen density.

This explains why high tumor burden requires higher BiTE doses, why antigen density thresholds exist [7], and provides a quantitative framework for optimizing immunotherapy dosing strategies.

#### Clinical implication

For patients with high blast burden (many high-density targets), current dosing may be insufficient. Our membrane-confined model with microvillus topology predicts that dose escalation is strictly necessary to overcome the antigen sink effect generated by the high local concentration of receptors at the synapse.

## 10 Biological interpretation and outlook

The generalized ternary framework developed here is intended as a theoretical complement to the original (**TBM**) model, tailored to membrane-confined systems with finite receptor copy numbers and realistic surface topology. It makes explicit the dependence of the ternary dose response and of reduced descriptors such as 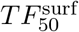 on antigen and effector receptor abundances. In the context of T cell-engaging therapies, this provides a principled way to interpret how differences in tumor-associated antigen density across cell lines or patients can reshape the window of ligand concentrations that yield substantial trimer formation at the immune synapse.

Several important biological processes are not yet included in the present formulation. These include CD3 and antigen internalization and recycling, spatial heterogeneity within the synapse beyond microvillus topology, and serial target engagement by a single T cell. The stochastic framework presented here can, in principle, accommodate such effects by adding sink and source terms to the master equation and by coupling multiple contact events over time. Furthermore, the trimer probability *P*_tri_(*t*) derived from our framework can be linked to functional readouts such as cytotoxicity by introducing a threshold number of productive trimers required for killing and integrating over the distribution of receptor densities in a cell population. These extensions will be addressed in future work and are particularly relevant for heterogeneity-aware models of BiTE dosing and hook-effect-constrained dose windows.

While we have illustrated the framework using the specific case of blinatumomab, the derivation relies only on general physical principles of membrane confinement and surface topology and is not limited to CD19/CD3 interactions. The accessibility factor (*η*) and microvillus concentration scaling apply equally to Chimeric Antigen Receptor (CAR) synapses, synthetic synNotch systems, and viral entry kinetics where receptors are confined to a contact interface. The finding that high receptor density can paradoxically increase *EC*_50_ in sequestration-dominated regimes is a general geometric property that should be considered in the design of any high-affinity bridging therapeutic.

## Acknowledgments

The authors thank colleagues for insightful discussions on three-body binding thermodynamics, membrane biophysics, and T cell engager pharmacology. Particular thanks to Allison Betts and John Gibbs both of *Takeda Development Center Americas, Inc*.

## Author Contributions

H.B. and K.P. initiated the project and developed the conceptual framework. H.B. derived all analytical and closed-form mathematical results and identified their physical and translational implications. K.P. contributed to the biophysical interpretation and clinical contextualization. H.B. and K.P. drafted the manuscript. All authors contributed revisions and additions and approved the submitted version

## Appendix A Dimensional analysis and units

Table 1 summarizes the units of the principal quantities appearing in the generalized ternary model.

**Table 1:** Units of selected quantities in the generalized ternary model.

| Quantity | Symbol | Units |
| --- | --- | --- |
| Contact area | $A_c$ | length<sup>2</sup> |
| Gap height | $h$ | length |
| Contact volume | $V_c$ | length<sup>3</sup> |
| Receptor counts | $N_A, N_C$ | molecules |
| Surface densities | $\sigma_A, \sigma_C$ | molecules per length<sup>2</sup> |
| Ligand concentration | $[B]_t$ | moles per length<sup>3</sup> |
| Three-dimensional $K_d$ | $K_{AB}, K_{BC}, K_T$ | moles per length<sup>3</sup> |
| Two-dimensional $K_d$ | $K_{AB}^{(2D)}, \dots$ | molecules per length<sup>2</sup> |

The effective receptor concentrations [*A*]^eff^, [*C*]^eff^ used in the effective concentration formulation are related to the surface densities by

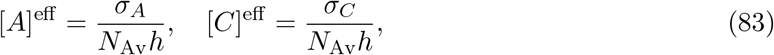

where *N*_Av_ is Avogadro’s number. This mapping preserves units and allows the direct use of three-dimensional dissociation constants. In the two-dimensional formulation, the dissociation constants can be defined by multiplying the three-dimensional constants by an appropriate length scale, such as the gap height *h*, as discussed in the main text.

## Appendix B

**Stochastic formulation and Poisson approximation**

For completeness, we sketch the derivation of the Poisson approximation for the trimer count distribution. Consider the reactions

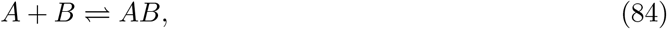

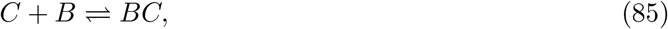

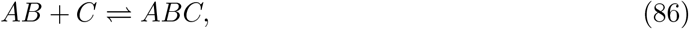

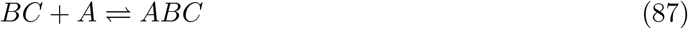

within the contact zone. The propensity functions for the forward and reverse reactions can be written in terms of the current counts of reactants and rate constants. For example, the formation of an AB dimer has a propensity

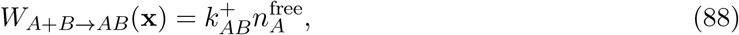

where 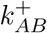 is an effective first-order rate constant that incorporates the ligand concentration [*B*]_loc_. Similar expressions hold for the other reactions.

The master equation for *P* (*n*_*AB*_, *n*_*BC*_, *n*_*ABC*_; *t*) is a linear system of ordinary differential equations. The first moment ⟨*n*_*ABC*_⟩ satisfies a closed equation in the mean-field approximation and in the long time limit approaches the deterministic equilibrium value.

When the mean trimer count is small compared to the available receptor copies, ⟨*n*_*ABC*_⟩ ≪ min(*N*_*A*_, *N*_*C*_), the distribution of *n*_*ABC*_ is well approximated by a Poisson distribution with mean ⟨*n*_*ABC*_⟩,

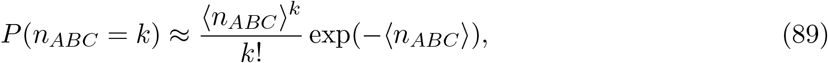

yielding

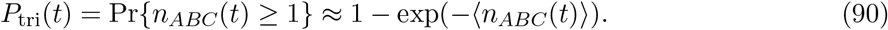

This approximation neglects the hard constraint that *n*_*ABC*_ ≤ min(*N*_*A*_, *N*_*C*_), but is accurate when trimer formation does not significantly deplete the receptor pools. This condition is typically satisfied in the formation regime and early stages of the balanced regime, but may break down when sequestration is very strong and a substantial fraction of receptors are bound. This approximation has been extensively used in models of receptor-mediated adhesion and bond clustering [2, 4] and provides a natural bridge from deterministic binding models to probabilistic statements about synapse formation and function.

## Appendix C

**Toy numerical illustration: effect of increasing antigen copy number**

To make the preceding discussion more concrete, we consider a minimal, dimensionless example comparing two antigen copy numbers, 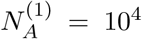 and 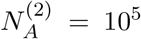, on an otherwise identical synapse.

### C.1 Setup and dimensionless parameters

We work within Branch A (effective concentration formulation) and focus on the target-dominant, noncooperative regime where the formation half point can be approximated as

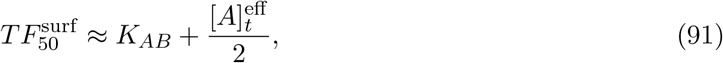

with 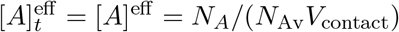. Rather than fixing absolute physical values of *V*_contact_ and *K*_*AB*_, we introduce dimensionless variables by choosing a reference scale for the target-side binding:

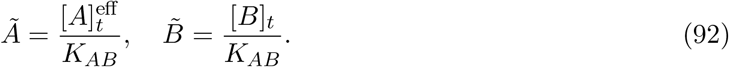

In these units, the approximate formation half point becomes

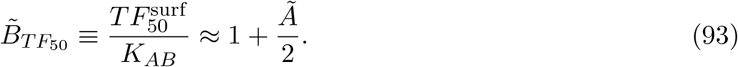

To keep the example simple, we assume that the effective concentration scales linearly with copy number via a constant geometric factor *κ*,

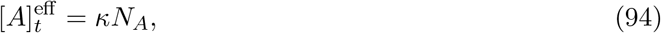

so that

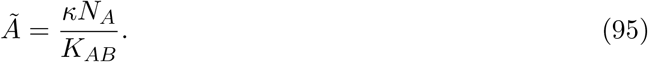

We choose units such that for the lower antigen density case 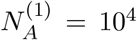, the dimensionless target load is modest,

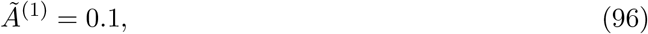

which fixes the ratio *κ/K*_*AB*_ via

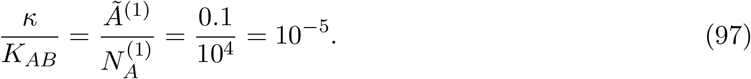

For the higher antigen copy number 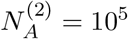, the dimensionless target load then becomes

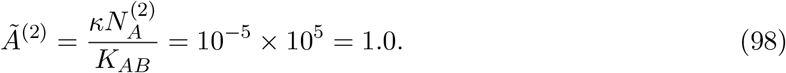

### C.2 Predicted shift in *TF*_50_

Substituting into the approximate expression (93), we obtain

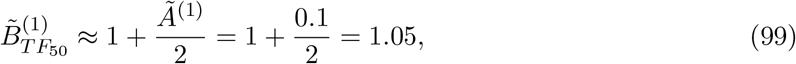

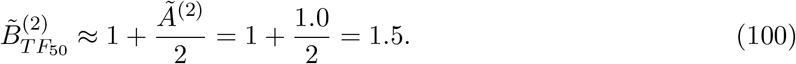

Thus, in this toy example, increasing the antigen copy number by one order of magnitude (10^4^ → 10^5^) produces:

- a tenfold increase in the dimensionless target load *Ã* (from 0.1 to 1.0), and
- an approximately 1.4-fold right shift in the formation half point, from 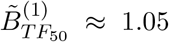 to 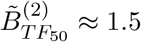.

At the same time, the maximal trimer level 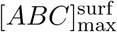 increases monotonically with 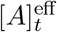, so the higher-TAA system not only supports more bridges per synapse but also requires a higher ligand dose (in these units) to reach half of its own maximal response.

### C.3 Interpretation

Even though the numbers above are purely illustrative, they capture the two key qualitative consequences of moving from a bulk description to a membrane-confined, finite-copy framework with microvillus topology:

1. At fixed geometry, the effective target concentration 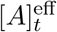 scales linearly with the copy number *N*_*A*_, so high-TAA synapses enter sequestration-dominated regimes at lower bulk doses than one would infer from a culture-scale bulk model.
2. When *TF*_50_ is defined relative to the maximum of the ternary curve, the combination of higher [*ABC*]_max_ and stronger dimer sinks causes 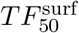 to shift to higher dosed [*B*]_*t*_ as *N*_*A*_ increases, even though the system is locally more favorable for trimer formation.

These trends provide a simple, quantitative illustration of why receptor copy number, synapse geometry, and microvillus topology must be made explicit when interpreting BiTE dose-response data and when comparing cell lines or patients with substantially different TAA levels.

#### Remark (Clarifying antigen dependence in (TBM) vs. the present model)

The classical bulk theory of Douglass [6] does include antigen abundance via the total concentration [*A*]_*t*_, and increasing [*A*]_*t*_ in their framework increases the maximal ternary complex [*ABC*]_max_. In the bulk formulation, however, [*A*]_*t*_ represents an effectively unbounded, spatially homogeneous reservoir of antigen that is not depleted locally by binding.

In contrast, our framework maps antigen abundance to a finite copy number of receptors confined to a synaptic contact area with microvillus topology. This introduces geometric and stoichiometric constraints through

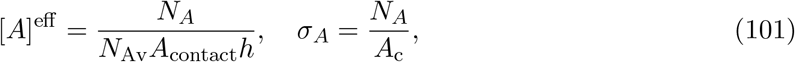

so that antigen can saturate, crowd, and sequester bridging ligand within the contact. As a consequence, increasing *N*_*A*_ raises [*ABC*]_max_ but can also right-shift the formation half point *TF*_50_ in sequestration-dominated regimes—an effect that does not arise from simply increasing [*A*]_*t*_ in the bulk theory. Thus our framework does not introduce antigen as a new variable, but rather embeds antigen into a finite membrane environment with realistic surface topology where geometry and copy number reshape the location of the ternary critical points.

